# Phosphate binding induced force-reversal occurs via slow backward cycling of cross-bridges

**DOI:** 10.1101/2020.08.20.259283

**Authors:** R Stehle

## Abstract

The release of inorganic phosphate (P_i_) from the cross-bridge is a pivotal step in the cross-bridge ATPase cycle leading to force generation. It is well known that P_i_ release and the force-generating step are reversible, thus increase of [P_i_] decreases isometric force by product inhibition and increases the rate constant *k*_TR_ of mechanically-induced force redevelopment due to the reversible redistribution of cross-bridges among non-force-generating and force-generating states. The experiments on cardiac myofibrils from guinea pig presented here show that increasing [P_i_] increases *k*_TR_ almost reciprocally to force, i.e., *k*_TR_ ≈ 1/force. To elucidate which cross-bridge models can explain the reciprocal *k*_TR_-force relation, simulations were performed for models varying in sequence and kinetics of 1) the P_i_ release-rebinding equilibrium, 2) the force-generating step and its reversal, and 3) the transitions limiting forward and backward cycling of cross-bridges between non-force-generating and force-generating states. Models consisting of fast reversible force generation before/after rapid P_i_ release-rebinding fail to describe the *k*_TR_–force relation observed in experiments. Models consistent with the experimental *k*_TR_-force relation have in common that P_i_ binding and/or force-reversal are/is intrinsically slow, i.e., either P_i_ binding or force-reversal or both limit backward cycling of cross-bridges from force-generating to non-force-generating states.

**STATEMENT OF SIGNIFICANCE:** Previous mechanical studies on muscle fibers, myofibrils and myosin interacting with actin revealed that force production associated to phosphate release from myosin’s active site presents a reversible process in the cross-bridge cycle. The correlation of this reversible process to the process(es) limiting kinetics of backward cycling from force-generating to non-force-generating states remained unclear.

Experimental data of cardiac myofibrils and model simulations show that the combined effects of [P_i_] on force and the rate constant of force redevelopment (*k*_TR_) are inconsistent with fast reversible force generation before/after rapid P_i_ release-rebinding. The minimum requirement in sequential models for successfully describing the experimentally observed nearly reciprocal change of force and *k*_TR_ is that either the P_i_ binding or the force-reversal step limit backward cycling.

## INTRODUCTION

Force generation of muscle is mechanically powered by the cross-bridge ATPase cycle. During this cycle, the cross-bridge passes numerous chemical and structural states which can be roughly grouped according their binding and force exerted on the actin filament into weakly-bound, non-force-generating and strongly-bound, force-generating states. The forward transition from non-force-generating to force-generating states is closely, albeit not necessarily directly, associated with release of inorganic phosphate (P_i_) from the active site of the cross-bridge. Elucidating the steps in the cycle that limit the transitions between non-force and force-generating states is fundamental for understanding the cross-bridge mechanism, the rate of contraction and relaxation and their targeted modulation by therapeutic interventions [reviewed in (1-8)].

Studies of muscle fibers contracting under high load or during stretch provide evidence that the P_i_ release is reversible, i.e. that cross-bridges can rebind P_i_ (9-13). However, different opinions exist whether the P_i_ is released before (14-19), along (20), after (21-23) or independent of (24) the step in the ATPase cycle that generates the force. The sequence of the P_i_ release and the force-generating step, also called the power stroke remains controversial (17, 22). Moreover, the kinetics of the P_i_ release-rebinding equilibrium as well as the nature of the rate-limiting processes controlling forward and backward transitions between non-force-generating and force-generating states poses unsolved problems (for reviews see (2, 5, 7, 8, 23, 25).

A standard technique for quantifying the rates limiting the transition between non-force-generating and force-generating states is to determine the kinetics of force redevelopment induced by switching from a transient period of active unloaded to active isometric contraction. The rate constant *k*_TR_ of this mechanically-induced force redevelopment represents the sum of apparent rate constants in the cross-bridge cycle limiting the transitions of cross-bridges between non-force-generating and force-generating states (26), reviewed in (2). There is consensus in several studies on slow and fast skeletal and cardiac muscles that P_i_ increases *k*_TR_ whereas it decreases force (7, 16, 27-37). The opposing effects of P_i_ on *k*_TR_ and force are in line with the reversibility of P_i_ release and the force-generating step in muscles contracting under mechanical load, i.e. rebinding of P_i_ shifts the equilibria from force-generating AM.ADP state(s) back to non-force-generating AM.ADP.P_i_ state(s) (9, 10, 13-15, 18, 38, 39). Though the sequence and kinetics of these reverse step(s) are unknown, the backward flux induced by P_i_ binding opens an additional pathway for redistribution of cross-bridges between non-force-generating and force-generating states. Without P_i_, this redistribution is solely determined by forward transitions in the ATPase cycle and *k*_TR_ = *f* + *g*, where *f* is the apparent rate constant of their forward transition into force-generating states and *g* the apparent rate constant of their forward transition into non-force-generating states. Increasing [P_i_] promotes rebinding of P_i_ to non-force states limited by the apparent rate constant *f* ^−^ that now contributes to *k*_TR_, i.e., *k*_TR_ = *f* + *g* + *f* ^−^, where *f* ^−^ is a function of the [P_i_] (7, 34, 40, 41).

Whereas in some models of the cross-bridge cycle the P_i_ release is indistinguishably connected to the step(s) limiting the transition of cross-bridges from non-force-generating to force states (20, 40, 42, 43), other models imply that P_i_ release occurs rapidly, independently from slower step(s) in the ATPase cycle (14-16, 18). Rapid increase in [P_i_] produced by flash photolysis of caged-P_i_ in muscle fibers induces a fast force decay with a rate constant *k*_Pi_ considerably higher than *k*_TR_ (14, 16). The higher value of *k*_Pi_ compared to *k*_TR_ and the dependence of *k*_Pi_ on [P_i_] were interpreted in terms of fast, reversible force-generation followed by rapid, reversible P_i_ release, i.e. by two subsequent fast reversible equilibrium, one for the power stroke and one for P_i_ release [(14, 16) reviewed in (8)]. In that model, rapid P_i_ binding and fast force-reversal determine the high *k*_Pi_ while redistribution among non-force and force-generating states is limited by other, slower transition(s) that determine the low *k*_TR_. However, this scenario has been questioned by studies exploring force kinetics upon rapid changes in [P_i_] in myofibrils [(33, 35) reviewed in (7)]. The thin myofibril bundles are suitable for studying force kinetics induced by rapid switching between two solutions of different [P_i_], enabling to change the [P_i_] in both directions. While rapid increases in [P_i_] induce fast force decays as in fibers, rapid decreases in [P_i_] induce slow force rises with *k*_-Pi_ similar to *k*_TR_. Furthermore, in cardiac myofibrils, the fast kinetics of force decay upon rapid increase in [P_i_] was attributed to sequential ‘give’ of sarcomeres (33), a phenomenon also observed during fast muscle relaxation (34, 44, 45). In contrast, the slow force rise upon rapid decrease in [P_i_] and the similarity of its rate constant *k*_-Pi_ with *k*_TR_ indicates that rapid perturbation of the P_i_ release-rebinding equilibrium induces a force response that is limited by similar slow processes as those limiting force redevelopment (7). Moreover, a novel regulatory, load-dependent mechanism involving organized conformational changes of the cross-bridge array on the thick filament between two states, OFF and ON, has been discovered (46). The possible modulation of cross-bridge turnover kinetics by this mechanism and novel insights into the structural cycle of myosin (4, 47, 48) further revive the question which steps in the cross-bridge cycle limit forward and backward transitions between non-force and force-generating states.

The present study explores the significance of the effects of [P_i_] on *k*_TR_ and force for the coupling of the P_i_ release-rebinding equilibrium and the transitions limiting redistribution between non-force-generating and force-generating states. Experimental data from cardiac myofibrils of guinea pig is compared with simulations of different models of the cross-bridge cycle differing in sequence and kinetics of three critical steps/events associated to myosin ATPase driven force-generation: reversible P_i_ release, reversible force-generating step and the event controlling the reversible transition of cross-bridges from weakly-bound to strongly-bound states. It is demonstrated that the interrelation of *k*_TR_ and force at various [P_i_], i.e. the P_i_-modulated *k*_TR_-force relation is a sensitive probe of the kinetic coupling between the P_i_-binding induced force-reversal and the rate-limiting backward transition *f* ^−^ and that either P_i_ binding or force-reversal or both have to be strongly coupled to *f* ^−^ for explaining the observed *k*_TR_-force relation.

Coupling strength becomes maximal when P_i_ binding and reversal of the force-generating step are merged with *f* ^−^ to a slow single step in a two state cross-bridge model (Fig. S1 in the Supporting Material), in which P_i_ alters *k*_TR_ and force in simple reciprocal relation, i.e. *k*_TR_ ≈ 1/force (see Eq. 1). Based on this limiting case, an empirical equation is given for defining coupling strength on a scale from +1 for maximum coupling and 0 for *k*_TR_ being independent on the [P_i_]. The P_i_-modulated *k*_TR_-force relation obtained from cardiac myofibrils from guinea pig yields a high coupling strength of +0.84 ± 0.08. Data from literature for other muscle preparations from cardiac, fast and slow skeletal muscle yields similar high coupling strengths indicating that the process of P_i_ induced force-reversal is directly coupled to the rate-limiting transition *f* ^−^.

## METHODS

### Myofibrillar preparation and solutions

Guinea pigs (Dunkin-Hartley, 450 – 750g) were anesthetized by 5 vol% isofluorane and sacrificed by decapitation. All use of animals and procedures in this study comply with the law for animal protection (TierSchG) transferred from EU guidelines and were reviewed and approved by the Official Animal Care and Use Committee (LANUV NRW, Az 84-02.05.20.13.080 and 84-02.05.50.15.029). After exsanguination of the animal body, the heart was excised and skinned strips from trabeculae prepared on the basis of (49). First, the blood was removed from the heart by brief (2-3 min) retrograde perfusion through the aorta at 37 °C with perfusion solution (132 mM NaCl, 5 mM KCl, 1 mM MgCl_2_, 10 mM TRIS, 5 mM EGTA, 1 mM sodium azide, 7 mM glucose, 2 mM DTT, pH 7.1). Then, the heart was placed in ice-cold perfusion solution without glucose and the left ventricular cavity opened by a cut in axial direction. Strips of 0.3 - 0.4 mm in diameter were dissected from the endocardial *trabeculae carneae* and pinned with microneedles on the sylgard surface of a chamber containing ice-cold skinning solution: 1% v/v Triton-X-100, 5 mM K-phosphate, 5 mM Na-azide, 3 mM Mg-acetate, 5 mM K_2_EGTA, 3 mM Na_2_ATP (incl. 3 mM MgCl_2_ and 6 mM KOH), 47 mM CrP, 2 mM DTT, 0.5 mM 4-(2-aminoethyl)benzenesulfonylfluoride HCl, 10 μM leupeptine, 10 μM antipaine, 5 mg/ml aprotinine (pH 7 at 0 °C). The pinned strips were incubated in the skinning solution at 0 °C for 4 h. Then, the skinning solution was replaced by storage solution (same composition as skinning solution but without triton) in which the skinned strips were stored at 4 °C for maximally 3 days. Myofibrils were prepared at the day of the mechanical experiment by homogenizing the skinned strip at 0 °C for 4-6 s at maximum speed with a blender (T10 Ultra-Turrax, IKA, Stauffen, Germany) and filtering the homogenate through polypropylene meshes (22 µm pore opening).

Standard activating buffer (pCa 4.5) for mechanical experiments contained 10 mM imidazole, 3 mM CaCl_2_K_4_EGTA, 1 mM Na_2_MgATP, 3 mM MgCl_2_, 47.7 mM Na_2_CrP, 2 mM DTT and different [P_i_] (pH 7.0 at 10 °C, µ = 0.178 M). Standard relaxation buffer (pCa 7) contained 3 mM K_4_Cl_2_EGTA instead of 3 mM CaCl_2_K_4_EGTA. Submaximal activating buffers pCa (5.88 5.03) were prepared by mixing standard activating and relaxing buffer at different ratios. Free calcium concentration [Ca^2+^] and pCa = −log [Ca^2+^]/M was calculated by a computer program (50). The [P_i_] in the buffers was measured by a phosphate assay kit (E-6646; Molecular Probes, Eugene, OR). P_i_ contamination in the standard activating buffer was 170 ± 20 µM (mean ± s.d.). Activating and relaxing buffers of lower [P_i_] (15 ± 5 µM P_i_) were produced by adding 1 mM methylguanosine and 0.5 units/ml purine nucleotide phosphorylase (PNP). Activating and relaxing buffers of higher [P_i_] were produced by adding phosphate buffer (30 % NaH_2_PO_4_^2-^ 70 % Na_2_HPO_4_^-^, pH 6.85). To hold ionic strength (µ) constant, [Na_2_CP] was reduced by 0.67 mM per 1 mM increase in [P_i_].

### Apparatus and technique to measure myofibrillar force redevelopment

The mechanical setup for measurements of myofibril force transients was described previously (34, 51). A droplet of myofibril suspension in storage solution was added to the thermostatically controlled (10 °C) chamber filled with relaxing solution. After sedimentation, a thin myofibril bundle was picked up from the chamber bottom at one end by a tungsten micro needle (# 5775, A-M Systems, Inc., Carlsborg, WA) and the other end moved with a manipulator to the tip of an atomic force cantilever (Nanoprobe^*©*^ FESP type, compliance: 0.2 0.4 µm/µN) that was coated with a mixture of 4 % nitrocellulose in amyl-acetate nitrocellulose and silicon adhesive (3140 RTV Coating, Dow Corning, Midland, USA). The free end of the bundle was pushed to the coating using a microneedle installed on a separate manipulator. Bundles used in experiments had diameters of 1.0 - 3.2 µm and slack lengths of 31 - 66 µm.

Slack sarcomere length (sSL) was 2.02 ± 0.11 µm (mean ± SD). Prior activation, bundles were stretched to 2.4 µm sarcomere length (SL). Signal conditioning for movement of actuators and acquisition of force and length signals were performed with a PCI6110-E device under self-written programs in LabView 4.0 (National Instruments, Austin, TX). During force recording, the myofibril was exposed to one of two laminar streams of solutions produced by a double channel theta-style capillary (TGC150-15, Clark Electromed. Instr., UK) and driven by gravitational pressure (30-35 cm H_2_O). Rapid Ca^2+^ activation and relaxation were induced by rapid solution change (52). The position of the flows was changed by rapid lateral movement of the capillary motored by a piezoactuator (P289.40, Physik Instr., Karlsruhe, Germany), effectively changing the solution at the bundle within 5-15 ms. Force redevelopment (*k*_TR_-measurement) was induced during Ca^2+^ activation. Rapid length changes were applied via the microneedle to the bundle using a piezoactuator (P602.1SL, Physik Instr.). To obtain the rate constant *k*_TR_ of force redevelopment, a single exponential function was fitted to the force transients (see Fig. 1B for fit examples) using a routine under LabView.

**FIGURE 1.**
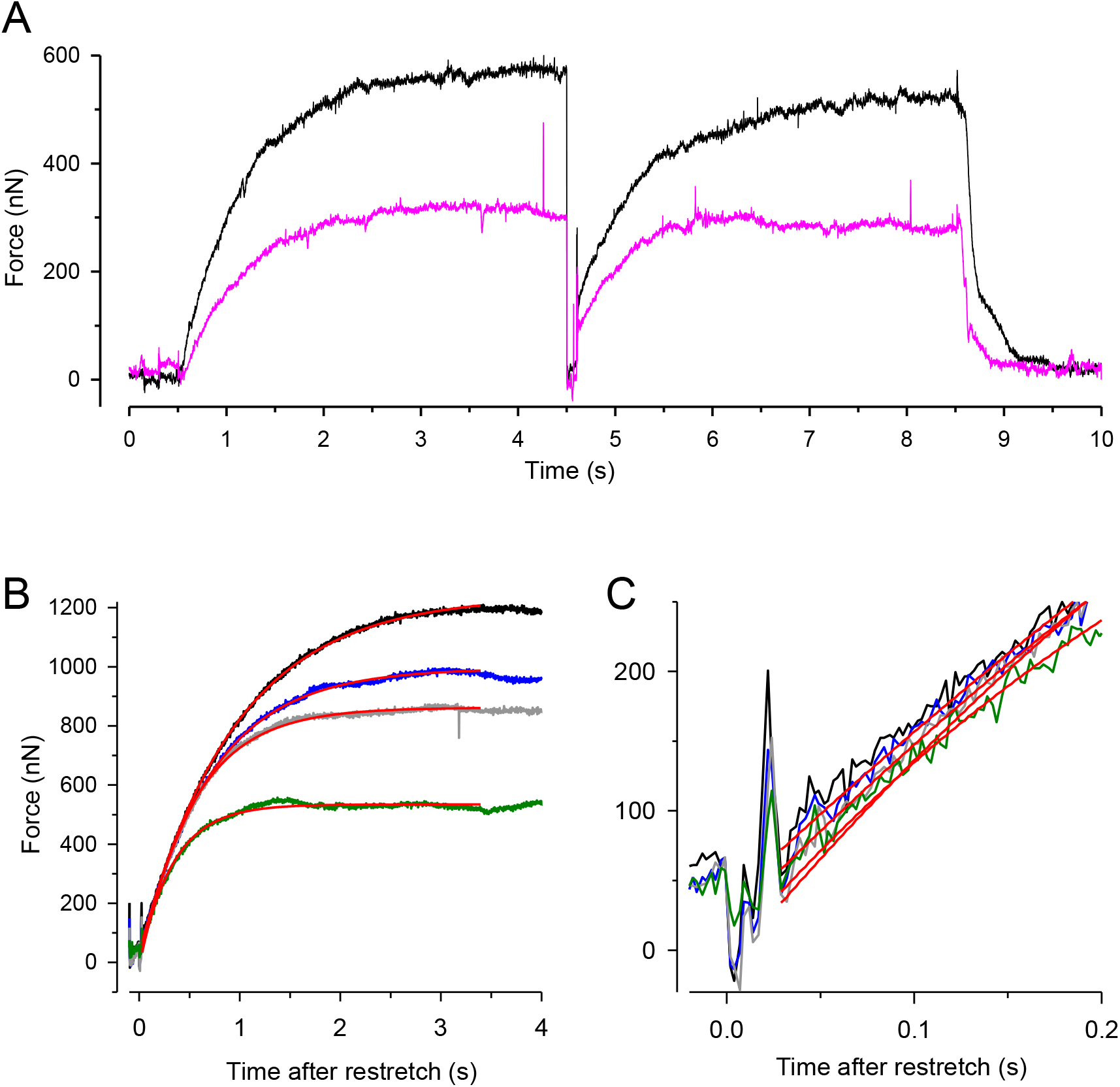
Experimental protocol and force redevelopment at different [P_i_]. (*A*) Typical full force transients obtained from a myofibril bundle (2.3 µm diameter, 66 µm length) at 0.17 ± 0.04 mM P_i_ (contaminant [P_i_] in standard buffer, black transient) and 20 mM P_i_ (pink transient). At *t* = 0.5 s, the bundle was activated by switching from relaxing solution (pCa 8) to activating solution (pCa 4.5). At *t* = 4.5 s, the bundle was slackened for 100 ms by 15 % of its length and then re-stretched to the original length to induce force redevelopment. At *t* = 8.5 s, the bundle was relaxed by switching back to relaxing solution. Force redevelopment after re-stretch mostly starts from a higher level than slack force like in this example. (*B*) Force redevelopment transients from a myofibril bundle (3.2 µm diameter, 47 µm length) at 1 mM P_i_ (*black*), 2.5 mM P_i_ (*blue*), 10 mM P_i_ (*grey*), and 20 mM P_i_ (*green*). Red lines are single exponentials fitted to transients yielding values for *k*_TR_ of 1.4 s^-1^ (1 mM P_i_), 1.7 s^-1^ (2.5 mM P_i_), 1.9 s^-1^ (10 mM P_i_), and 2.9 s^-1^ (20 mM P_i_). In this experiment, force redevelopments started close to slack force enabling the comparison of their initial force rises which are similar in slope as shown in (*C*).

### Model simulations and definition of coupling strength

Model simulations were performed under Berkeley Madonna 8.3.18 (for rate constants see Table 1).

**TABLE 1.**
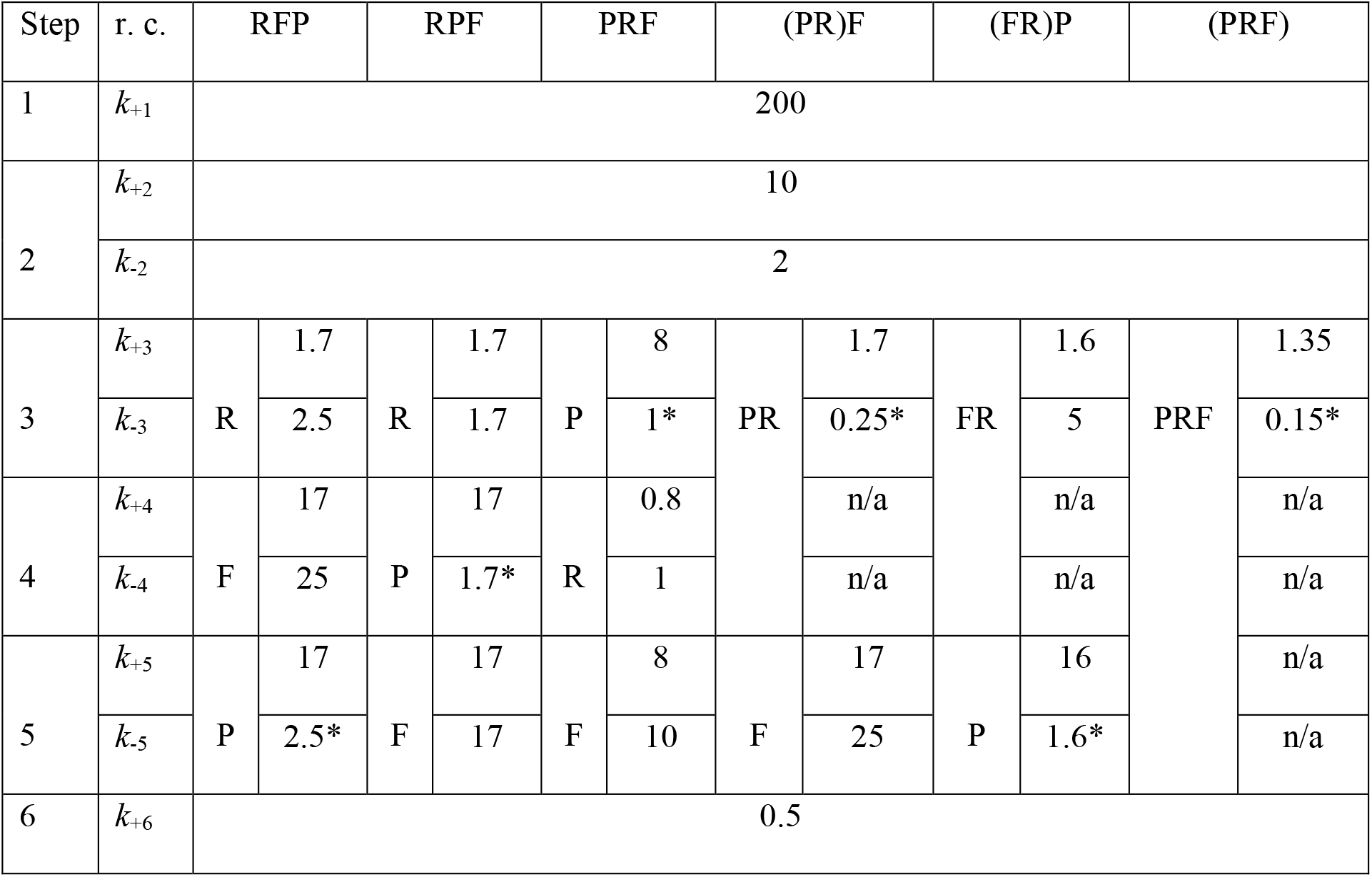
Rate constants (r. c.) used for simulations of 6 different models of the cross-bridge cycle. Step 1 and step 6 are assumed to be irreversible (reverse rate constants = 0), steps 2 – 5 are reversible equilibria. All models are equal in step 1 (ATP binding), step 2 (ATP hydrolysis), and step 6 (load dependent isomerization including ADP release). Models differ in attribution of step 3 - 5 to the reversible equilibria R, F, and P where R presents the equilibrium of the rate-limiting forward and backward transitions *f* and *f* ^*–*^, F presents the equilibrium of the force-generating step and its reversal, and P presents the equilibrium of P_i_ release-rebinding. Name of models indicate the sequence of R, F and P in the cycle. Parentheses in names mean that P or F or both are merged with R to single slow equilibrium which results in omitted steps (rate constant: n/a) and less than 6 steps in that models. Unit of rate constants is s^-1^ except for the second order rate constant of P_i_ rebinding, *k*_B_ [mM^-1^s^-1^]*. Final values of rate constants in models were set according following criteria: 1) When P or F are fast equilibria separate from R, the rate constants of P or F are 10-fold those of R. 2) The forward rate constant of R is set to match the observed *k*_TR_ at low [P_i_]. In the special case of the PRF model, the sum of forward and reverse rate constants of R is set to match the observed *k*_TR_ at low [P_i_]. 3) The reverse rate constants of R, F and P are set to match the force reduction at high [P_i_].

As described in more detail in the Supporting Material (Appendix), an indicator, the coupling strength (*CS*) was defined to quantify the coupling between P_i_ binding induced force reduction and the rate-limiting backward transition *f* ^−^ in the cross-bridge ATPase cycle. *CS* was defined on scales from 0 for no coupling to +1 for maximum coupling and from 0 to]-1 for maximum inverse coupling.

Briefly, by definition, *CS* = 0 refers to the case when *k*_TR_ remains constant (*k*_TR_ = *const*.), independent of the force changes by P_i_.

Maximum positive coupling (*CS* = +1) is reached in case of a two state model when P_i_ rebinding, force reversal and *f* ^−^ are merged to a single step (Fig. S1), i.e. when all three steps are the same. In this case altering [P_i_] modulates *k*_TR_ directly inversely to force by

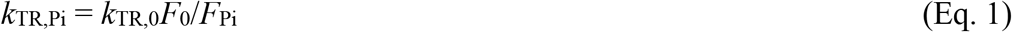

where *k*_TR,Pi_ is *k*_TR_ and *F*_Pi_ is the force at a given [P_i_]. *k*_TR,0_ is *k*_TR_ and *F*_0_ is the force at basal [P_i_].

Positive *CS* in the interval [0, +1] is described by

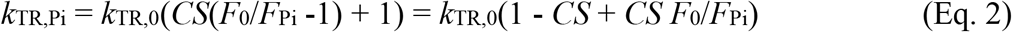

Negative coupling strength in the interval [-1, 0] is described by

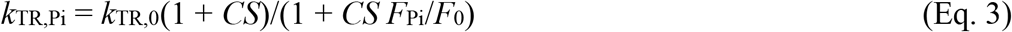

Eq. 2 and Eq. 3 can be fit to *k*_TR_-force data of muscle preparations obtained at various [P_i_] to derive the *CS* from experimental data. For comprehensive understanding how *k*_TR_-force relations depend on *CS*, see family of curves plotted in Fig. S2.

## RESULTS

### Experimental protocol and characteristics of force redevelopment at different [P_i_]

Fig. 1 shows force transients of guinea pig cardiac myofibrils at 10 °C, pCa 4.5 and different [P_i_]. The experimental protocol is illustrated by full force recordings in Fig. 1A. The myofibril bundle is exposed to the flow of relaxing solution (pCa 8) and then active force development is initiated by rapidly switching to the flow of activating solution (pCa 4.5) consisting of the same [P_i_] as the relaxing solution. During steady state Ca^2+^ activation, force redevelopment (*k*_TR_-measurement) is induced by a slack-restretch manoeuvre. After redevelopment of force, the bundle is relaxed by switching back from activating to relaxing solution. Then, the next activation-*k*_TR_-measurement-relaxation cycle is performed at the next [P_i_].

Increasing [P_i_] reduces the isometric force and the time for reaching the force plateau (Fig. 1A and 1B). Force transients are fitted by single exponential functions (red lines in Fig. 1B and 2C) to determine the rate constant of tension redevelopment *k*_TR_. Increasing [P_i_] from 1 mM up to 20 mM P_i_ reduces force down to ∼50 % and increases *k*_TR_ up to ∼2.1-fold (Fig. 1B) while hardly changing (by less than 15 %) the initial slope of the force redevelopment (Fig. 1C). A constant initial slope is expected when the rate constant of an exponential function changes reciprocally to its amplitude, reflecting the case of maximum possible rate-modulation of *k*_TR_ (see Eq. 1).

**FIGURE 2.**
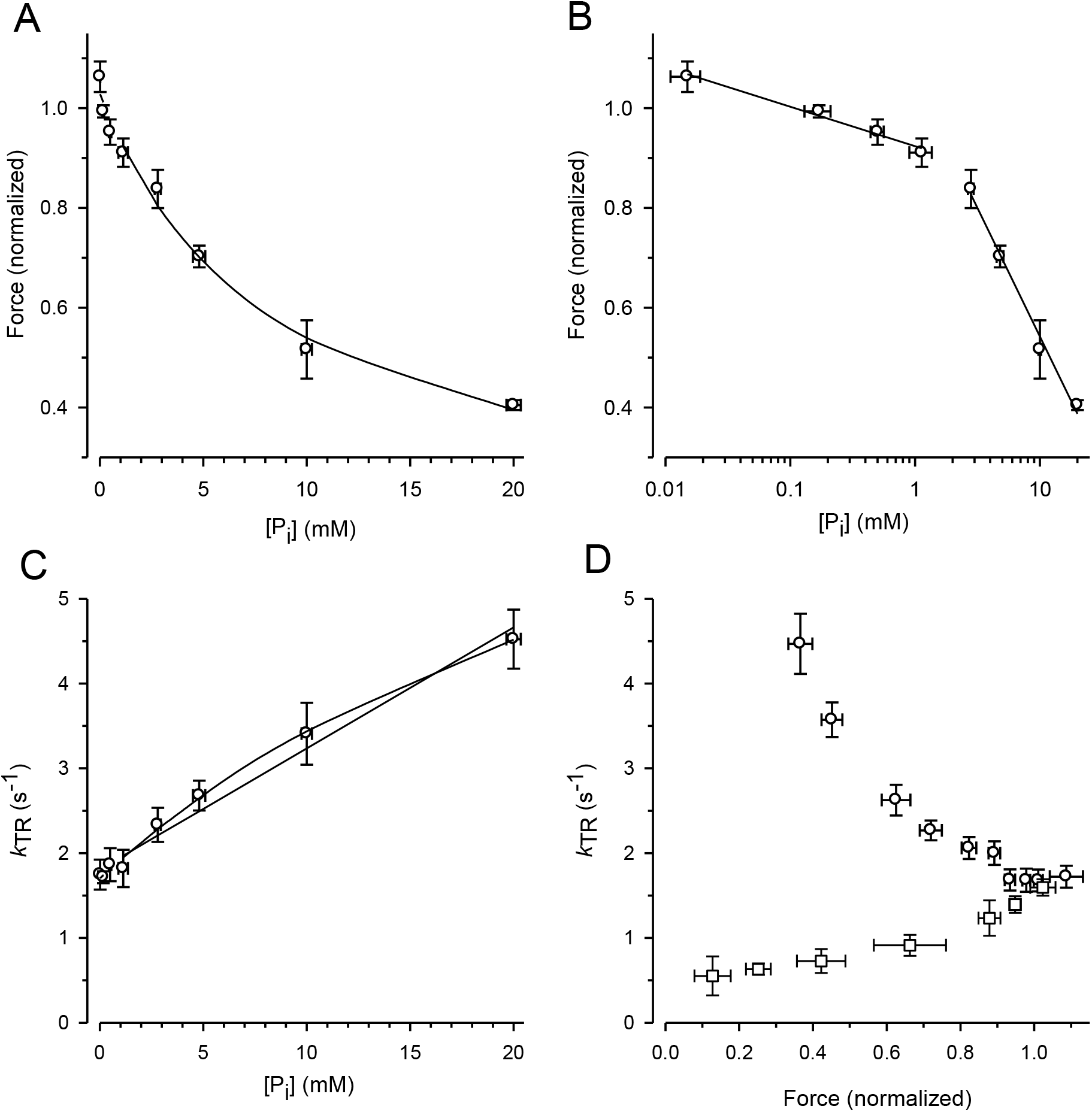
Alteration of force and *k*_TR_ by [P_i_] and comparison of *k*_TR_-force relations resulting from varying [P_i_] and [Ca^2+^]. For each myofibril, force data was normalized to force measured in standard activating solution (0.17 mM P_i_, pCa 4.5). (*A*) Force-[P_i_] relation. The line present the hyperbolic function fitted to the data yielding *F*_0Pi_ = 1.03 ± 0.02, *P*_*i*50_ = 8.4 ± 2.1 mM and *F*_∞Pi_ = 0.13 ± 0.09. (*B*) Force-log [P_i_] relation. Lines indicate linear regression lines with slopes of −0.08 per decade increase of [P_i_] at low [P_i_] (≤ 1 mM P_i_) and −0.51 at high [P_i_] (≥ 2.5 mM P_i_). (*C*) *k*_TR_-[P_i_] relation and analysis of its curvature. The lines present linear (*k*_0Pi_ = 1.81 ± 0.07 s^-1^, slope = 0.143 ± 0.008 s^-1^/mM P_i_) or hyperbolic fit functions (*k*_0Pi_ = 1.70 ± 0.04 s^-1^, *P*_*i*50_ = 33 ± 9 mM and *k*_∞Pi_ = 9.2 ± 1.3 s^-1^) to the data. (*D*) Relations of *k*_TR_ versus force altered either by changing the [P_i_] between 0.015 mM and 20 mM (*circles*) at full Ca^2+^ activation (pCa 4.5) or by changing the pCa between 4.5 and 5.88 (*squares*) at constant [P_i_] of 0.17 mM. Data are means ± SEM

### Dependence of force and *k*_TR_ on the [P_i_]

To exploit force reduction and rate modulation of *k*_TR_ over a broad range of [P_i_], force redevelopment transients were recorded from 19 myofibril bundles at variable [P_i_] ranging from 0.015 mM to 20 mM P_i_. The force values of the transients were then normalized to the mean force produced by the myofibril bundle in the standard activating solution which contained a contaminant [P_i_] of 0.17 mM. Fig. 2A shows the relation of the normalized active force on [P_i_]. Force is already reduced by sub-millimolar [P_i_]. Fitting the force-[P_i_] relation by a hyperbolic function yields three parameters: the fit value at zero [P_i_] (*F*_0Pi_), the [P_i_] for half-maximum hyperbolic change (*P*_*i*50_) and the final value at infinite [P_i_] (*F*_∞Pi_). The fitted *F*_∞Pi_ of 0.13 ± 0.09 might point to a small active force component that cannot be completely reversed by P_i_. Plotting force on a logarithmic scale of [P_i_] reveals a bi-linear relation with a 6-fold less steep decrease of force per decade increase of [P_i_] for the data ≤ 1 mM P_i_ than for the data ≥ 2.5 mM P_i_ (Fig. 2B).

Fig. 2C shows the increase of the *k*_TR_–data with the [P_i_] which can be fitted with a linear and hyperbolic function. If a process apart from P_i_ binding limits backward transition from force to non-force states (*f* ^−^), then *k*_TR_ saturates at high [P_i_] and the hyperbolic *k*_TR_-[P_i_] relation is expected. In contrast, if *f* ^−^ refers to the rate constant of P_i_ binding, the linear increase of *k*_TR_ with [P_i_] is expected. There is some weak curvature in the *k*_TR_-[P_i_] relation and the hyperbola fits better to the data than the linear (lines in Fig. 2C). However, considering the error of the data and the additional variable implemented in the hyperbolic compared to the linear function, the data does not discriminate one of the above scenarios.

The effects of [P_i_] on *k*_TR_ and force are integrated in the form of the *k*_TR_-force relation in which the *k*_TR_ values are paired with the relative force obtained in the same transient (Fig. 2D, circles). Regarded from the right to the left, according the direction of increasing [P_i_], decrease of force is accompanied with an increase of *k*_TR_ whereby the increment of *k*_TR_ per decrement of force becomes higher at lower force.

To exploit whether *k*_TR_ simply increases due to the lower isometric force, force was reduced by reducing the [Ca^2+^] in the standard activating solution without added P_i_. Force transients of 8 myofibrils were recorded at full and partial Ca^2+^ activation, force of each transient normalized to force at full Ca^2+^ activation (pCa 4.5, 0.17 mM P_i_) and the *k*_TR_ value paired with the normalized force of the same transient and plotted in Fig. 2D (square symbols). In line with several previous studies, Ca^2+^ modulates *k*_TR_ in same direction as force (26, 28, 30, 32, 53-56), i.e. in opposite direction to the P_i_–modulated *k*_TR_-force relation.

### Quantification of coupling strength from [P_i_]-modulated *k*_TR_-force data of cardiac myofibrils

To quantify the *CS* from the experiments varying the [P_i_], the individual *k*_TR_ obtained from each transient was paired with the normalized force value of the same transient and the data pairs plotted in the *k*_TR,Pi_–force relation shown in Fig. 3. The symbols present the values of 144 force transients obtained from 19 myofibrils at different [P_i_] (indicated by the different symbols (or color in the online version). The line presents the best fit of Eq. 2 to the data yielding *CS* = 0.84 ± 0.08 and *k*_TR,0_ = 1.73 ± 0.07 s^-1^. The latter reflects the *k*_TR_-value of the fit curve at the unity force in the standard solution, i.e. at 0.17 mM P_i_.

**FIGURE 3.**
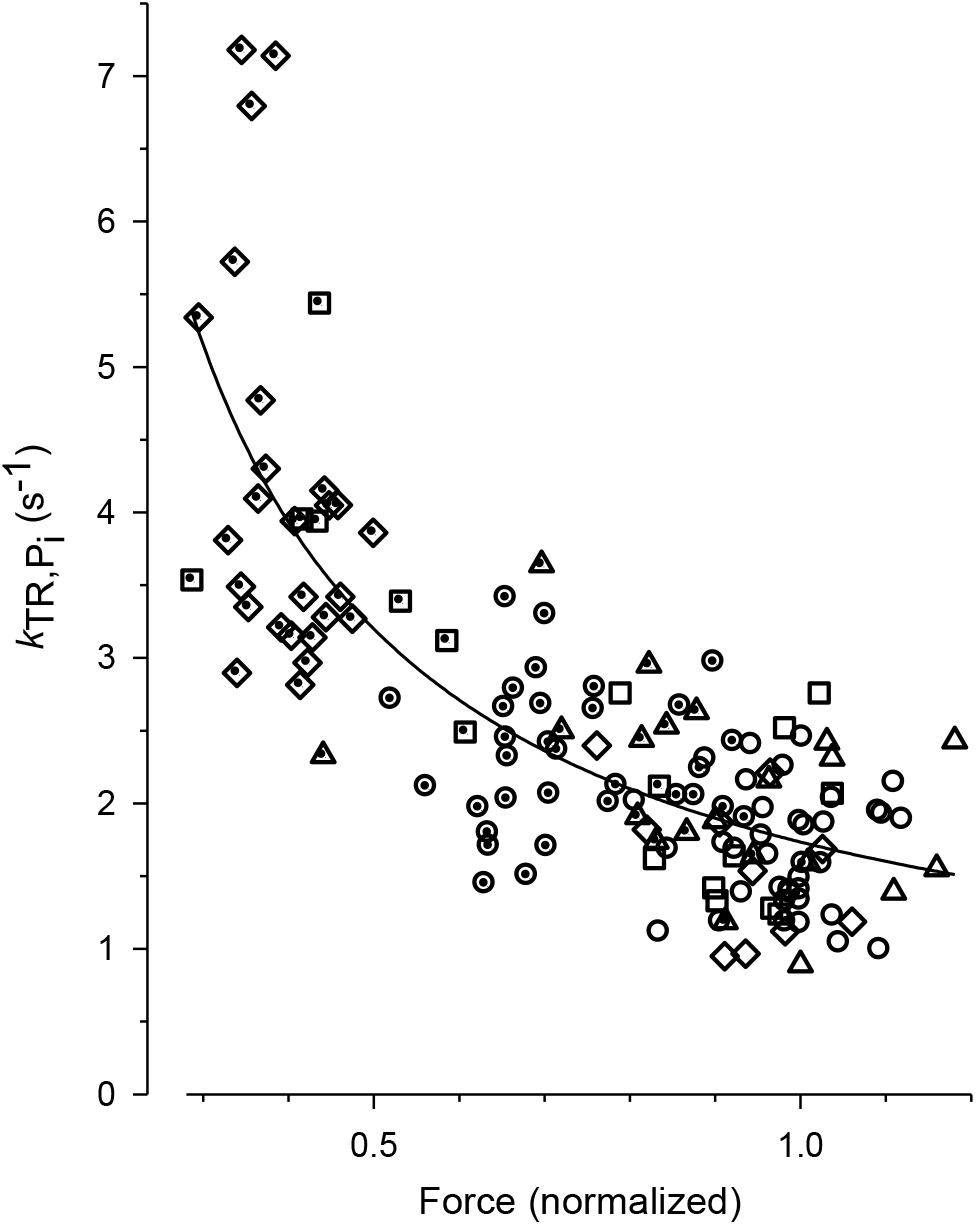
Fit of coupling strength function (Eq. 2) to *k*_TR_-force data. Force data is normalized to mean force of each myofibril at 0.17 mM P_i_ (contaminant P_i_ in standard solution). In 3 bundles, [P_i_] was further reduced by the P_i_ scavenger PNP resulting in a [P_i_] of 0.015 mM (*triangles*, data of 9 transients). *Circles*: data obtained at 0.17 mM P_i_ (36 transients), s*quares*: 0.5 mM P_i_ (10 transients), *diamonds*: 1 mM P_i_ (10 transients), *dotted triangles*: 2.5 mM P_i_ (12 transients), *dotted circles*: 5 mM P_i_ (33 transients), *dotted squares*: 10 mM P_i_ (8 transients), *dotted diamonds*: 20 mM P_i_ (26 transients). The best fit of Eq. 2 (line) to the pooled *k*_TR_-force data yields the fit coefficients *CS* = 0.84 ± 0.08 and *k*_TR,0_ = 1.73 ± 0.07 s^-1^ (mean ± s.d).

### Rate modulation of *k*_TR_ by [P_i_] and coupling strength depends on cross-bridge model

To test the compatibility of the high *CS* obtained in the myofibril experiments with models of the cross-bridge cycle, models differing in sequence and kinetics of three critical events determining the reversible transition into force-generating states were tested for their [P_i_]-dependent modulation of force and *k*_TR_. The three critical events were defined as reversible equilibria, an equilibrium abbreviated by R for the rate-limiting forward and backward transition (*f* and *f* ^−^), an equilibrium abbreviated by F for the force-generating step and its reversal, and an equilibrium abbreviated by P for P_i_ release-rebinding. The equilibria R, F and P were embed in various models of the cross-bridge cycle using same rate constants for ATP binding (step 1), ATP hydrolysis (step 2), and the load dependent ADP release (step 6) but different sequences of R, F and P (step 3 - 5) and different association of P or F with R. To simulate the scenarios of F or P or both presenting the rate-limiting forward-backward transition, they were merged with R to a single equilibrium. The different models and their rate constants are summarized in Table 1.

For each model, force redevelopment transients were simulated at various [P_i_]. Each transient was produced by first calculating the steady-state distribution of states during unloaded shortening for which the forward rate constant of step 6 was set to high value for unloaded shortening (*k*’_+6_ = 50 s^-1^) and then switching it at *t* = 0 to low value for isometric contraction (*k*_+6_ = 0.5 s^-1^). The resulting transients simulating the isometric force redevelopments were then fitted by the same type of single exponential function used for the transients in real myofibril experiments for obtaining the *k*_TR_-values and force of each model.

The force amplitudes in each model were normalized to the force amplitude at 0.17 mM P_i_ and the normalized force plotted against the [P_i_] (Fig. 4A) and the log [P_i_] (Fig. 4B) together with the relations obtained in the myofibril experiments. The curvature of the myofibril force-[P_i_] relation can be largely described by the models (PFR) and (PR)F in which P_i_ release/rebinding limits forward/backward transition into/from force-generating states, except at the highest [P_i_] of 20 mM P_i_ where both models overestimate the observed force reduction. Similar curvature of the force-[P_i_] relation is predicted by the RPF-model with the sequence of the rate-limiting transition controlling rapid P_i_ release triggering a fast force-generating step. Models in which force is generated before rapid P_i_ release predict increased curvatures, regardless whether F is coupled to R in the (FR)P model or F is a fast step behind R in the RFP model. Lowest curvature is predicted by the PRF model in which rapid P_i_ release antecedes the rate-limiting transition. Nevertheless, all models recapitulate the basic feature of force reduction over a large range of [P_i_], making it difficult to exclude certain models based on force-[P_i_] and force-log [P_i_] relations.

**FIGURE 4.**
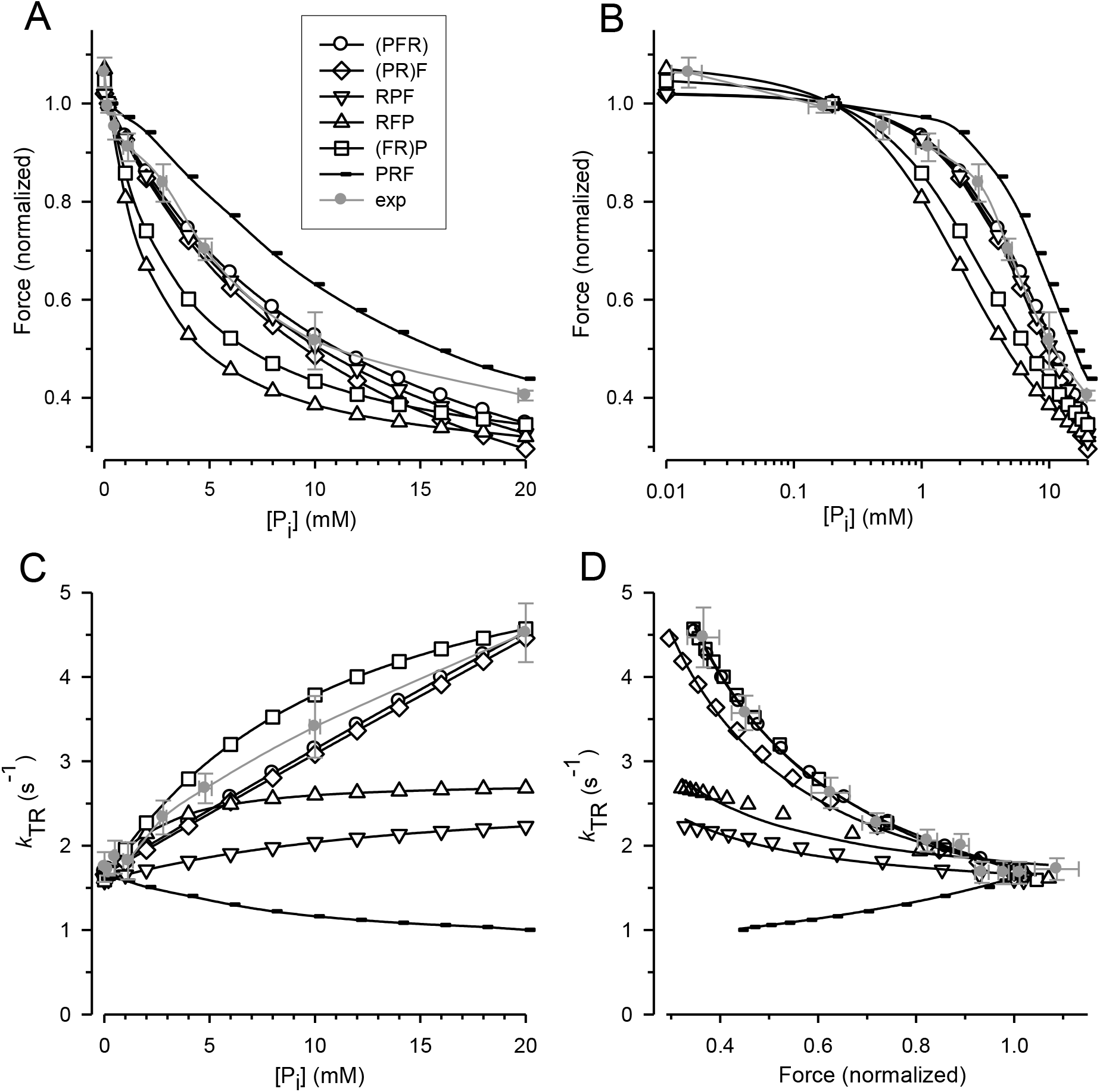
Relation of force and *k*_TR_ on [P_i_] predicted by cross-bridge models differing in sequence and kinetics of reversible equilibria for P_i_ release and force-generating step. (*A*) Force-[P_i_] relations. (*B*) Force-log [P_i_] relations. (*C*) *k*_TR_-[P_i_] relations. (*D*) *k*_TR_-force relations. *Filled grey circles* and error bars (red in online version) represent the experimental data replotted from Figure 3. Black symbols indicate relations calculated for the different models: *Open circles* refer to the (PFR) model in which P_i_ release, force-generating step and rate-limiting transition *f* are merged to a single slow step. *Squares* refer to the (FR)P model in which the force-generating step presents the rate-limiting transition *f* followed by faster P_i_ release. *Diamonds* refer to the (PR)F model in which the P_i_ release presents the rate-limiting transition *f* followed by a faster force-generating step. *Tip-up triangles* refer to the RFP-model with the sequence: 1. rate-limiting transition *f*, 2. fast force-generating step, 3. rapid P_i_ release. *Tip-down triangles* refer to the RPF model with the sequence: 1. rate-limiting transition *f*, 2. rapid P_i_ release, 3. fast force-generating step. *Small symbols* present the PRF model with the sequence: 1. rapid P_i_ release, 2. rate-limiting transition *f*, 3. fast force-generating step. Lines in subfigures (*A*-*C*) are spline curves. Lines in (*D*) represent best fits of Eq. 2 to each model except RPF model to which Eq. 3 was fitted. Model-dependent *CS*: 0.86 ± 0.02 (PFR), 0.90 ± 0.02 (FR)P, 0.67 ± 0.02 (PR)F, 0.26 ± 0.03 RFP, 0.19 ± 0.03 RPF, and −0.52 ± 0.01 PRF.

Fig. 4C shows that the *k*_TR_-[P_i_] relation depends on the type of the model. In the (PFR) and (PR)F models, in which the backward transition *f* ^*–*^ is directly limited by P_i_ rebinding, *k*_TR_ increases steeply, linearly with the [P_i_] (Fig. 4C). Also steep but curved increase of *k*_TR_ with [P_i_] is observed when force generation and its reversal is coupled to the rate-limiting transitions prior rapid P_i_ release-rebinding as in the (FR)P model. The *k*_TR_-[P_i_] relation of cardiac myofibrils scales in between the linear relations of the (PFR) and (PR)F models and the curved relation of the (FR)P model and would be consistent with these three models. In contrast, models in which F and P are fast, reversible equilibria results in less steep *k*_TR_-[P_i_] relations than observed in experiments. In even stronger contrast, the PRF model in which rapid P_i_ release-rebinding occurs before the rate-limiting transitions *f* and *f* ^-^ results in a declining *k*_TR_-[P_i_] relation.

To obtain the coupling strength (*CS*), Eq. 2 was fitted to the *k*_TR_-force data simulated for each model (Fig. 4D). Models in which P_i_ release and rebinding are merged with rate-limiting transitions yield high *CS* of 0.86 for the (PRF) model and 0.90 for the (PR)F model (Fig. 4D). The *CS* of these models does not reach the maximum value of 1.0 as in case of a two-state model because the ATP cleavage step also slightly participates in limiting redistribution of cross-bridges between non-force and force-generating states. The *CS* of both models are in agreement with the *CS* of 0.84 ± 0.08 obtained in real experiments (Fig. 3). Reasonable high *CS* of 0.67 is also obtained in the (FR)P model where fast P_i_ release-rebinding occurs after rate-limiting, reversible force generation. However, when both, force-generating step and P_i_ release are assumed to be fast reversible equilibria apart from the rate-limiting transition *f* and *f* ^*–*^, *CS* becomes low. Neither the scenario of a fast reversible force-generating step triggering fast reversible P_i_ release (RFP model, *CS* = 0.26) nor the scenario of fast reversible P_i_ release step triggering fast reversible force-generation (RPF model, *CS* = 0.19) can account for the high *CS* observed in experiments. Finally, the PRF model in which rapid reversible P_i_ release antecedes the rate-limiting transition to force-generating states results in negative *CS* of −0.51.

## DISCUSSION

### Implications of the force-[P_i_] relation for mechanism of force generation

The asymptote of the hyperbolic fit to the force-[P_i_] relation yields an estimate of the relative active force remaining at infinite [P_i_] (*F*_∞Pi_). A value of *F*_∞Pi_ greater than 0 indicates that saturating [P_i_] cannot fully inhibit active force. The remaining force could either result from limited reversibility of P_i_ release or from force-generating AM.ADP.P_i_ states (15, 16, 57). Because all models simulated in Fig. 4 assume P_i_ release to be fully reversible, only the force-[P_i_] relations of the RFP and (FR)P models in which force is generated before P_i_ release do not approach zero force at infinite [P_i_] (Fig. 4A). In all other models, *F*_∞Pi_ = 0. The value of *F*_∞Pi_ = 0.13 ± 0.09 in this study might point to a low active force at infinite [P_i_]. Tesi and coworkers explored force-[P_i_] relation of rabbit psoas myofibrils up to 70 mM P_i_ and obtained an asymptote value of 0.07 (35, 36). Considering the low *F*_∞Pi_ and limited [P_i_] range, the force-[P_i_] relations of myofibrils from cardiac and fast skeletal muscle provide no definite answer for or against force-producing AM.ADP.P_i_ state(s). Noteworthy, skinned fiber preparations of fast skeletal muscle (16, 37, 58-61) exhibit much higher asymptote values than myofibrils of this muscle type (35, 36) whereas fibers (62), myocytes (27, 29) and myofibrils (33) from cardiac muscle yield similar, low asymptote values. The reason for the less efficient force reduction by P_i_ in the fast muscle fiber preparation is unknown. However, force modulation by P_i_ will be underestimated when P_i_ accumulates and cross-bridge strain is not uniformly distributed in the preparation (63, 64). These technical limitations might be of particular importance at high ATPase activity and large size of preparations (62). Regardless of these technical limitations, myofibrils from slow rabbit soleus muscle (36) yield higher asymptote values than myofibrils from fast rabbit psoas muscle (36) and cardiac myofibrils (this study). One possible explanation would be that P_i_ release is less reversible in slow skeletal than in other muscles. The difference is not simply related to the myosin heavy chain (MHC) isoform since rabbit soleus myofibrils (36) and guinea pig cardiac myofibrils (this study) contain slow β-MHC (65).

The significance of the shape of the force-log [P_i_] relation for the force-generating mechanism has been pointed out in previous studies on skeletal and cardiac muscle preparations (27, 35, 66). Based on reduced models consisting of reversible equilibria for the force-generating step and the P_i_ release not implemented in a full cross-bridge cycle, a nearly mono-linear force-log [P_i_] relation is expected when force generation and P_i_ release occur in the same step (one-step model), whereas a sigmoidal relation is expected when force generation occurs before P_i_ release (two-step model). In a previous study on cardiac muscle, a mono-linear and a sigmoid function were fitted to force-log [P_i_] data obtained from skinned rat myocytes (27). Both fits do not match the experimental data and a bi-linear fit function might fit better to that data in (27). Tesi et al. explored rabbit psoas myofibrils at 5 °C and were the first fitting their data by using such type of bi-linear function (35). The slopes of the two linear lines in (35) were −0.07 and −0.40 which are similar to the −0.08 and −0.51 in this study (see Fig. 2B) indicating close similarity of force-log [P_i_] relations in fast skeletal and cardiac myofibrils. The force-log [P_i_] relation obtained for skinned rat myocytes was interpreted to be no evidence for the two-step model (27) while the data of myofibrils from fast skeletal muscle was interpreted to be in rough agreement with the two-step model (35). The slopes of the first linear at low [P_i_] up to 1 mM P_i_ for fast skeletal myofibrils (35) and cardiac myofibrils (Fig. 2B, this study) appear to be higher than the slope of the sigmoidal relation for a simple two-step model. However, a larger strain distribution of force-generating cross-bridges will also increase the slope in this [P_i_] range (67). It is noteworthy, that only myofibrils enable to explore the of force-log [P_i_] relation in the sub-millimolar range because the ATPase activity and long lateral diffusion distance of skinned fibers result in lateral [P_i_] gradients of 1-2 mM P_i_ in skinned fibers (64). Regarding the similarity of the force-log [P_i_] relations in fast skeletal and cardiac myofibrils and the different interpretations of force-log [P_i_] relations in previous studies of the two muscle types (27, 35), the shape of the force-log [P_i_] relation is not a very helpful criterion for deciding between the one-step and the two-step model.

One progress in this study compared to previous studies analyzing force-log [P_i_] relations (27, 35) is the use of models of the full cross-bridge cycle instead of one-step or two-step models without integration of these steps in a cycle. Interestingly, when regarding the full [P_i_] range, the force-[P_i_] and force-log [P_i_] relations simulated for different full cycle models appear to be very similar to each other (see Fig. 4A and 4B). Moreover, at low [P_i_], the full cycle models yield some contradictory predictions to the prediction of ‘isolated step’ models. Thus, the (PFR) model that integrates the one-step model in the cycle yields a flatter force-log [P_i_] relation than the RFP model that integrates the two-step model. The full cycle model simulation reveals that the overall shape of the force-log [P_i_] relation is rather insensitive to coupling mechanism between force generation and P_i_ release and that the slope of force-log [P_i_] relation at low [P_i_] depends more complex on the coupling mechanism than previously thought.

### Implications of the *k*_TR_-[P_i_] relation for the mechanism of force generation

Addition of 10 mM P_i_ in skinned cardiac myocytes from human donor hearts increases *k*_TR_ 2.4-fold (30) similar to the 2.0-fold increase of *k*_TR_ by 10 mM P_i_ here observed in cardiac myofibrils from guinea pig. In rat skinned cardiac myocytes, addition of 10 mM P_i_ increases *k*_TR_ 3.8-fold (29), and in cardiomyocytes from human, pig and mouse, 1.5-fold, 1,6-fold and 2.9-fold, respectively (28). The stronger P_i_ effects in mouse and rat compared to human, pig and guinea pig heart might be partly related to the fast α-MHC isoform expressed in murine and rat ventricles compared to the slow isoform expressed in human, pig and guinea pig (65). However, the myosin isoform unlikely is the only determinant of the *k*_TR_-[P_i_] relation as myofibril and fiber preparations from slow skeletal muscle exhibit no change in *k*_TR_ (35-37). The insensitivity of *k*_TR_ to [P_i_] in slow skeletal muscle could point to an incomplete reversibility of P_i_ release or different force-generating mechanism in slow skeletal muscle (7).

The model simulations in this study reveal that slope and the curvature of the *k*_TR_-[P_i_] relation is sensitive to the force-generating mechanism. The slope is strongly positive for models in which P_i_ binding induced force-reversal is slow and limits *f* ^−^, it is flat for models in which P_i_ binding induced force-reversal is a fast process and it becomes negative for models in which P_i_ release-rebinding is a fast equilibrium before the rate-limiting transitions *f* and *f* ^−^ in the cycle. The *k*_TR_-[P_i_] relation is linear when P_i_ binding directly limits the transition of cross-bridges from force-generating to non-force-generating states and curved downward when P_i_ binding is fast so that another process gets rate-limiting for this transition at high [P_i_]. Despite of its clear predictions for the mechanism, no clear picture emerges from the curvature of the *k*_TR_-[P_i_] relations reported in literature. Downward curved *k*_TR_-[P_i_] relations were reported for skinned fast (32, 37) and slow (37) muscle fibers, a slightly downward curved relation for myofibrils from rabbit psoas (35) and an upward curvature for skinned rat cardiac myocytes (29). The slightly downward curved *k*_TR_-[P_i_] relation obtained here with cardiac myofibrils can be fitted by a hyperbola as well as by a linear function (see Fig. 2C) corroborating the difficulty to decide on the mechanism from the shape of the *k*_TR_-[P_i_] relation. Hence, an alternative, quantitative approach is required to analyze the modulation of *k*_TR_ and force by [P_i_]. This was done here by analyzing the combined modulation of *k*_TR_ and force by [P_i_], i.e. by analyzing the P_i_-modulated *k*_TR_–force relation in terms of *CS*.

### Implications of the *k*_TR_-force relation and the *CS* for the mechanism of force generation

The present study depicts the first quantitative analysis of the coupling strength *CS* between the process of P_i_ binding induced force reduction and the transition limiting backward cycling represented by the rate constant *f* ^-^. Empirical equations are given in Methods and Supporting Material to derive the *CS* from *k*_TR_-force relations observed in real experiments and theoretical model simulations. Applying the equations to model simulations reveal that starting from the case of maximum *CS* = 1 for a two-state model, in bottom-up direction towards more extended models, *CS* remains high as long as the reversible equilibrium for P_i_ release or the reversible equilibrium for force-generation or both equilibria remain bound to the rate-limiting forward-backward transitions (see models (PFR), (PR)F, (FR)P in Fig. 4D) and becomes low when both of them are separated from this transitions (see models RFP and RPF). On a scale from +1 for maximum positive over zero for no coupling to −1 for maximum inverse coupling, the *k*_TR_-force relation of cardiac myofibrils yields a *CS* of 0.84 close to 1, consistent with the (PFR), (PR)F, and (FR)P models but inconsistent with the RFP and RPF models.

Due to the linear scaling of positive *CS* defined by Eq. 2 and shown in Fig. S2A, the *CS* reflects the ratio of the relative increase of *k*_TR_ per relative decrease of force induced by a certain increase of [P_i_]. Thus, the *CS* of 0.84 in this study means that in average, *k*_TR_ increases by 0.84-times the reduction of force, e.g. *k*_TR_ increases 0.84 x 2-fold = 1.68-fold when force is reduced 2-fold to ½ of its initial value. Albeit, to the best of the author’s knowledge, no previous study regarded this ratio, numerous studies reported *k*_TR_ and force values containing this information. Addition of 10 mM P_i_ in rat skinned cardiac myocytes increases *k*_TR_ 3.8-fold while it reduces force 3-fold (29). Addition of 10 mM P_i_ in skinned cardiac myocytes from human donor hearts increases *k*_TR_ 2.4-fold while it reduces force 2.5-fold (30) similar to the 2.0-fold increase of *k*_TR_ and 2.0-fold reduction of force here observed in cardiac myofibrils from guinea pig. In fast skeletal muscle fibers, addition of ≥ 10 mM P_i_ doubled *k*_TR_ and halved force (37) and in fast skeletal myofibrils, force was already halved by addition of 5 mM P_i_ along with 3-fold increase of *k*_TR_ (35, 36). In summary, the data of cardiac and fast skeletal muscle show the tendency of P_i_ changing *k*_TR_ roughly reciprocally to force as expected for the maximum *CS* of 1. However, the insensitivity of *k*_TR_ to P_i_ in fibers (37) and myofibrils (36) from slow skeletal muscle would yield low *CS*, even when considering the lower effects of P_i_ on force in this muscle type.

The high *CS* in fast skeletal and cardiac muscle might underlie either slow rebinding of P_i_, i.e., low *k*_B_ as in (PFR) and (PR)F models, or slow reversal of the power stroke as in (RFP) and (FR)P models. Low *k*_B_ is not necessarily in conflict with the classic theory of rapid, diffusion-limited ligand binding because P_i_ is released and rebound through so-called backdoor(s) which might limit dissociation and binding kinetics of P_i_ (4, 68, 69). Model simulations also reveal high *CS* for the (FR)P model in which force-generation is coupled to rate-limiting transitions prior rapid reversible P_i_ release. Thus, high *CS* does not exclude the classical rapid, diffusion-limited reaction for P_i_ binding. However, in this case, the rate-limiting backward transition to non-force-generating states expressed by *f* ^−^ has to be coupled to the reversal of the force-generating step.

The observed high *CS* is in conflict with classical two-step mechanisms consisting of an intermediate fast reversible force generating step followed by rapid reversible P_i_ release where both equilibria are faster than the transitions controlling forward and backward flux of cross-bridges into and back from force-generating states (8, 14-16, 18). This scenario is reflected by the RFP model which yields *CS* = 0.26, i.e. a value much lower than the *CS* of 0.84 observed in cardiac myofibril experiments (Fig. 4D). Using the rate constants given in the classical studies favoring the RFP model for rabbit psoas muscle at 10 °C (14, 16) also results in low *CS* of 0.28 (Fig. S3). *CS* even gets lower when the sequence of the two fast reversible equilibria is permuted and force generation occurs after P_i_ release as shown by the RFP model in Fig. 4D yielding *CS* = 0.19.

Noteworthy, *CS* be ‘rescued’ by making two modifications in the RFP model. Firstly, by lowering *k*_B_ whereby slow P_i_ binding gets back rate-limiting for backward cycling of cross-bridges while P_i_ release can be kept fast, i.e. the rate constant *k*_P_ can be kept high. Thereby, rate modulation of *k*_TR_ by [P_i_] gets restored but with the side effect of loss of force modulation by [P_i_] due to the high *k*_P_/*k*_B_ ratio, i.e. the high equilibrium constant *K*_P_ of reversible P_i_ release. The latter effect can be restored by secondly lowering the equilibrium constant of the reversible force-generating step F (*K*_F_). The low *K*_F_ and high *K*_P_ result in very low occupancy of the post-power stroke, pre-P_i_ release state (force-generating AM_F_.ADP.P_i_ state). The final result is a model in which the low occupancy of the initial state of the P_i_ release (AM_F_.ADP.P_i_ state) limits the rate of P_i_ release even though the intrinsic rate constant of P_i_ release (*k*_P_) is high and *k*_TR_ is modulated by [P_i_] via slow rebinding of P_i_ (low *k*_B_). In this type of model, forward transition of cross-bridges to force-generating states is limited by a lowly occupied post-power stroke, pre-P_i_ release state from which P_i_ is rapidly released. Even this type of model consists of fast power stroke and subsequent rapid P_i_ release in forward direction, it appears to be essential for obtaining high *CS* that either P_i_ binding or force-reversal in backward direction is slow and coupled to *f* ^−^.

### Model simplifications, assumptions and limitations

The models simulated here only regard the ‘horizontal’ steps of cross-bridge turnover while the ‘vertical’ steps for actin binding of cross-bridges are neglected. The basic conclusions from the present analysis also apply for models involving actin binding when the rate constants of the horizontal steps are regarded as apparent rate constants reflecting the rate constant of the horizontal step times the actin bound occupancy of its initial state.

The major assumptions for deriving the *CS* by Eq. 2 are that 1) force redevelopment kinetics reflects cross-bridge turnover kinetics, 2) the rate constant of force development presents the sum of rate constants limiting redistribution among non-force- and force-generating states, and 3) there is only one step in the cycle generating the force, and 4) there is only one main sequential pathway of rate-limiting transition *f*, power stroke and P_i_ release.

The first assumption implies that cross-bridges cycle independently of each other with same kinetic properties. Recently, it has been shown that the cross-bridges on the thick filament shifts from a structural OFF state to an ON state when switching from low load shortening to high load contraction (46, 47). It is presently unknown whether the redistribution of thick filament states participates in limiting cross-bridge turnover kinetics. The effect of P_i_ on this redistribution still needs to be explored.

The second assumption implies that rate constants are independent of time and do not change during force redevelopment. This was put recently in question by Kawai based on the argument that cross-bridges have to cycle many times because their step size is much less than the distance of filament sliding during force redevelopment (70). However, this argument does not apply in case of multiple interactions (24) or reversible detachment-reattachment processes (71) of force-generating cross-bridges. A prediction of Kawai’s model (70) is that *k*_TR_ inversely relates to tension cost given by the ratio of isometric tension per ATPase. Tension cost is independent of Ca^2+^ activation (26) and increases ∼2-fold with increasing [P_i_] to 30 mM in fast skeletal and cardiac muscle (59, 72, 73). The effect on tension cost is ∼5-fold or 7.5-fold too low to account for the ∼10-fold or ∼15-fold difference observed in *k*_TR_ when force is reduced by altering Ca^2+^ and P_i_ in cardiac myofibrils (see Fig. 2D) or fast skeletal muscle fibers (32), respectively. The classical interpretation of *k*_TR_ provides a simple explanation for these large differences and opposite changes of *k*_TR_ resulting from force reduction via Ca^2+^ and P_i_ by decreasing *f*_app_ and increasing *f*_app_^−^, respectively (reviewed in (7)).

The third assumption is only an approximation as single myosin force measurements indicate the presence of a second force-generating step associated to ADP release (74). A reversible force-generating step beside P_i_ release would lower the observed modulation of the *k*_TR_-force relation, decrease the *CS* value derived from this relation and thus lead to an underestimation of the coupling between P_i_ binding induced force-reversal and *f* ^*–*^. The high *CS* derived from the myofibril experiments indicates that the second force generating step does not contribute very much to the derived *CS*, either because this step is irreversible or minor in size (74).

Contrary to the fourth assumption, some studies suggest branched kinetic pathways (75-77), multiple structural pathways of P_i_ release through different backdoors (68) or binding of P_i_ to different chemical cross-bridge states (78). The analysis in the present study does not account for these alternatives. Simplification to a single sequential pathway is a usual requirement for working out the main kinetic and structural cross-bridge cycle.

## Conclusions

Increase of [P_i_] strongly increases the rate constant of force redevelopment *k*_TR_ in cardiac myofibrils as previously observed in cardiac and skeletal muscle fibers and myofibrils from skeletal muscle. Acceleration of *k*_TR_ by P_i_ does not simply result from the force reduction by P_i_ because reducing force by lowering the [Ca^2+^] causes the well-known opposed effect of decreasing *k*_TR_. Correlating the effects of P_i_ on *k*_TR_ and force in a *k*_TR_-force relation provides a quantitative test of the coupling between the P_i_ binding step, the reversal of the force-generating step and the transition limiting the backward flux of cross-bridges to non-force-generating states expressed by the rate constant *f* ^*–*^. Strong [P_i_]-dependent rate modulation of *k*_TR_ requires *f* ^*–*^ to be limited by the P_i_ binding step and/or the reverse step of force-generation. Simulations of *k*_TR_-force relations using models in which P_i_ release-rebinding is a rapid equilibrium prior or subsequent to a rapid reversible force-generating step yield low rate modulation of *k*_TR_ by [P_i_], in contrast to strong modulation observed in experiments in cardiac and skeletal muscle preparations.

The finding of slow backward kinetics of P_i_ binding induced force-reversal indicated by the *k*_TR_-force relation complements the previous finding of slow kinetics of the myofibril force rise induced by rapidly changing from high initial [P_i_] to low final [P_i_] (7, 33, 35). At low final [P_i_], P_i_ binding becomes negligible and the rate constant *k*_–Pi_ of this slow force rise merely reflects forward kinetics. The similarity of *k*_–Pi_ and *k*_TR_ indicated that force generation associated to P_i_ release cannot be separated from the rate-limiting transition *f* (7, 33, 35). The strong rate modulation of *k*_TR_ by P_i_ indicates that P_i_ rebinding induced force reversal cannot be separated from *f* ^*–*^. In combination, this indicates that force generation associated to P_i_ release and their reversal cannot be separated from the rate-limiting transitions *f* and *f* ^−^. The natures of these rate-limiting transitions still have to be defined in the structural model of the myosin motor ATPase cycle (48).

## AUTHOR CONTRIBUTIONS

RS designed the study, performed and analyzed the myofibril experiments and model simulations and wrote the manuscript.

## ACKNOWLEDGEMENT

This work was supported by Köln Fortune (Faculty of Medicine, Cologne) to RS.

## SUPPORTING CITATIONS

**References (79 – 81)** appear in the Supporting Material.

## REFERENCES

1. Geeves, M. A. 2016. Review: The ATPase mechanism of myosin and actomyosin. Biopolymers 105:483–491.

2. Gordon, A. M., E. Homsher, and M. Regnier. 2000. Regulation of contraction in striated muscle. Physiol Rev 80:853–924.

3. Hinken, A. C., and R. J. Solaro. 2007. A dominant role of cardiac molecular motors in the intrinsic regulation of ventricular ejection and relaxation. Physiology (Bethesda) 22:73–80.

4. Houdusse, A., and H. L. Sweeney. 2016. How Myosin Generates Force on Actin Filaments. Trends Biochem Sci 41:989–997.

5. Mansson, A., D. Rassier, and G. Tsiavaliaris. 2015. Poorly understood aspects of striated muscle contraction. Biomed Res Int 2015:245154.

6. Stehle, R., and B. Iorga. 2010. Kinetics of cardiac sarcomeric processes and rate-limiting steps in contraction and relaxation. J Mol Cell Cardiol 48:843–850.

7. Stehle, R., and C. Tesi. 2017. Kinetic coupling of phosphate release, force generation and rate-limiting steps in the cross-bridge cycle. J Muscle Res Cell Motil 38:275–289.

8. Takagi, Y., H. Shuman, and Y. E. Goldman. 2004. Coupling between phosphate release and force generation in muscle actomyosin. Philos Trans R Soc Lond B Biol Sci 359:1913–1920.

9. Hibberd, M. G., J. A. Dantzig, D. R. Trentham, and Y. E. Goldman. 1985. Phosphate release and force generation in skeletal muscle fibers. Science 228:1317–1319.

10. Hibberd, M. G., M. R. Webb, Y. E. Goldman, and D. R. Trentham. 1985. Oxygen exchange between phosphate and water accompanies calcium-regulated ATPase activity of skinned fibers from rabbit skeletal muscle. J Biol Chem 260:3496–3500.

11. Mannherz, H. G. 1970. On the reversibility of the biochemical reactions of muscular contraction during the absorption of negative work. FEBS Lett 10:233–236.

12. Ulbrich, M., and J. C. Ruegg. 1971. Stretch induced formation of ATP-32P in glycerinated fibres of insect flight muscle. Experientia 27:45–46.

13. Webb, M. R., M. G. Hibberd, Y. E. Goldman, and D. R. Trentham. 1986. Oxygen exchange between Pi in the medium and water during ATP hydrolysis mediated by skinned fibers from rabbit skeletal muscle. Evidence for Pi binding to a force-generating state. J Biol Chem 261:15557–15564.

14. Dantzig, J. A., Y. E. Goldman, N. C. Millar, J. Lacktis, and E. Homsher. 1992. Reversal of the cross-bridge force-generating transition by photogeneration of phosphate in rabbit psoas muscle fibres. J. Physiol. 451:247–278.

15. Kawai, M., and H. R. Halvorson. 1991. Two step mechanism of phosphate release and the mechanism of force generation in chemically skinned fibers of rabbit psoas muscle. Biophys J 59:329–342.

16. Millar, N. C., and E. Homsher. 1990. The effect of phosphate and calcium on force generation in glycerinated rabbit skeletal muscle fibers. A steady-state and transient kinetic study. J Biol Chem 265:20234–20240.

17. Muretta, J. M., J. A. Rohde, D. O. Johnsrud, S. Cornea, and D. D. Thomas. 2015. Direct real-time detection of the structural and biochemical events in the myosin power stroke. Proc Natl Acad Sci USA 112:14272–14277.

18. Ranatunga, K. W. 1999. Effects of inorganic phosphate on endothermic force generation in muscle. Proceedings Biological sciences 266:1381–1385.

19. Woody, M. S., D. A. Winkelmann, M. Capitanio, E. M. Ostap, and Y. E. Goldman. 2019. Single molecule mechanics resolves the earliest events in force generation by cardiac myosin. Elife 8.

20. Eisenberg, E., T. L. Hill, and Y. Chen. 1980. Cross-bridge model of muscle contraction. Quantitative analysis. Biophys J 29:195–227.

21. Davis, J. S., and M. E. Rodgers. 1995. Indirect coupling of phosphate release to de novo tension generation during muscle contraction. Proc Natl Acad Sci USA 92:10482–10486.

22. Llinas, P., T. Isabet, L. Song, V. Ropars, B. Zong, H. Benisty, … A. Houdusse. 2015. How actin initiates the motor activity of Myosin. Dev Cell 33:401–412.

23. Rahman, M. A., M. Usaj, D. E. Rassier, and A. Mansson. 2018. Blebbistatin Effects Expose Hidden Secrets in the Force-Generating Cycle of Actin and Myosin. Biophys J 115:386–397.

24. Caremani, M., L. Melli, M. Dolfi, V. Lombardi, and M. Linari. 2013. The working stroke of the myosin II motor in muscle is not tightly coupled to release of orthophosphate from its active site. J Physiol 591:5187–5205.

25. Geeves, M. A., and K. C. Holmes. 2005. The molecular mechanism of muscle contraction. Adv Protein Chem 71:161–193.

26. Brenner, B. 1988. Effect of Ca^2+^ on cross-bridge turnover kinetics in skinned single rabbit psoas fibers: implications for regulation of muscle contraction. Proc Natl Acad Sci USA 85:3265–3269.

27. Araujo, A., and J. W. Walker. 1996. Phosphate release and force generation in cardiac myocytes investigated with caged phosphate and caged calcium. Biophys J 70:2316–2326.

28. Edes, I. F., D. Czuriga, G. Csanyi, S. Chlopicki, F. A. Recchia, A. Borbely, … Z. Papp. 2007. Rate of tension redevelopment is not modulated by sarcomere length in permeabilized human, murine, and porcine cardiomyocytes. Am J Physiol Regul Integr Comp Physiol 293:R20–29.

29. Hinken, A. C., and K. S. McDonald. 2004. Inorganic phosphate speeds loaded shortening in rat skinned cardiac myocytes. American journal of physiology Cell physiology 287:C500–507.

30. Papp, Z., J. van der Velden, A. Borbely, I. Edes, and G. J. M. Stienen. 2014. Altered myocardial force generation in end-stage human heart failure. ESC Heart Fail 1:160–165.

31. Regnier, M., and E. Homsher. 1998. The effect of ATP analogs on posthydrolytic and force development steps in skinned skeletal muscle fibers. Biophys J 74:3059–3071.

32. Regnier, M., C. Morris, and E. Homsher. 1995. Regulation of the cross-bridge transition from a weakly to strongly bound state in skinned rabbit muscle fibers. Am J Physiol 269:C1532–1539.

33. Stehle, R. 2017. Force Responses and Sarcomere Dynamics of Cardiac Myofibrils Induced by Rapid Changes in [Pi]. Biophys J 112:356–367.

34. Stehle, R., M. Kruger, and G. Pfitzer. 2002. Force kinetics and individual sarcomere dynamics in cardiac myofibrils after rapid Ca^2+^ changes. Biophys J 83:2152–2161.

35. Tesi, C., F. Colomo, S. Nencini, N. Piroddi, and C. Poggesi. 2000. The effect of inorganic phosphate on force generation in single myofibrils from rabbit skeletal muscle. Biophys J 78:3081–3092.

36. Tesi, C., F. Colomo, N. Piroddi, and C. Poggesi. 2002. Characterization of the cross-bridge force-generating step using inorganic phosphate and BDM in myofibrils from rabbit skeletal muscles. J. Physiol. 541:187–199.

37. Wahr, P. A., H. C. Cantor, and J. M. Metzger. 1997. Nucleotide-dependent contractile properties of Ca^2+^-activated fast and slow skeletal muscle fibers. Biophys J 72:822–834.

38. Mansfield, C., T. G. West, N. A. Curtin, and M. A. Ferenczi. 2012. Stretch of contracting cardiac muscle abruptly decreases the rate of phosphate release at high and low calcium. J Biol Chem 287:25696–25705.

39. Woodward, M., and E. P. Debold. 2018. Acidosis and Phosphate Directly Reduce Myosin’s Force-Generating Capacity Through Distinct Molecular Mechanisms. Frontiers in physiology 9:862.

40. A, M. A., R. S. A, H. Q. R, A. F. M, A. A. M, A. H. F, and M. T. I. 2014. Women Health in Saudi Arabia: A review of non-communicable diseases and their risk factors. Pak J Med Sci 30:422–431.

41. Palmer, S., and J. C. Kentish. 1998. Roles of Ca^2+^ and crossbridge kinetics in determining the maximum rates of Ca^2+^ activation and relaxation in rat and guinea pig skinned trabeculae. Circ Res 83:179–186.

42. Huxley, A. F. 1957. Muscle structure and theories of contraction. Prog Biophys Biophys Chem 7:255–318.

43. Lymn, R. W., and E. W. Taylor. 1971. Mechanism of adenosine triphosphate hydrolysis by actomyosin. Biochemistry 10:4617–4624.

44. Flitney, F. W., and D. G. Hirst. 1978. Cross-bridge detachment and sarcomere ‘give’ during stretch of active frog’s muscle. J Physiol 276:449–465.

45. Huxley, A. F., and R. M. Simmons. 1970. Rapid ‘give’ and the tension ‘shoulder’ in the relaxation of frog muscle fibres. J Physiol 210:32P–33P.

46. Linari, M., E. Brunello, M. Reconditi, L. Fusi, M. Caremani, T. Narayanan, … M. Irving. 2015. Force generation by skeletal muscle is controlled by mechanosensing in myosin filaments. Nature 528:276–279.

47. Irving, M. 2017. Regulation of Contraction by the Thick Filaments inSkeletal Muscle. Biophys J 113:2579–2594.

48. Robert-Paganin, J., O. Pylypenko, C. Kikuti, H. L. Sweeney, and A. Houdusse. 2020. Force Generation by Myosin Motors: A Structural Perspective. Chem Rev 120:5–35.

49. Linke, W. A., M. L. Bartoo, and G. H. Pollack. 1993. Spontaneous sarcomeric oscillations at intermediate activation levels in single isolated cardiac myofibrils. Circ Res 73:724–734.

50. Fabiato, A., and F. Fabiato. 1979. Calculator programs for computing the composition of the solutions containing multiple metals and ligands used for experiments in skinned muscle cells. Journal de physiologie 75:463–505.

51. Stehle, R., M. Kruger, P. Scherer, K. Brixius, R. H. Schwinger, and G. Pfitzer. 2002. Isometric force kinetics upon rapid activation and relaxation of mouse, guinea pig and human heart muscle studied on the subcellular myofibrillar level. Basic Res Cardiol 97 Suppl 1:I127–135.

52. Colomo, F., S. Nencini, N. Piroddi, C. Poggesi, and C. Tesi. 1998. Calcium dependence of the apparent rate of force generation in single striated muscle myofibrils activated by rapid solution changes. Adv. Exp. Med. Biol. 453:373–381.

53. Kreutziger, K. L., N. Piroddi, B. Scellini, C. Tesi, C. Poggesi, and M. Regnier. 2008. Thin filament Ca^2+^ binding properties and regulatory unit interactions alter kinetics of tension development and relaxation in rabbit skeletal muscle. The Journal of physiology 586:3683–3700.

54. Norman, C., J. A. Rall, S. B. Tikunova, and J. P. Davis. 2007. Modulation of the rate of cardiac muscle contraction by troponin C constructs with various calcium binding affinities. American journal of physiology Heart and circulatory physiology 293:H2580–2587.

55. Sweeney, H. L., and J. T. Stull. 1990. Alteration of cross-bridge kinetics by myosin light chain phosphorylation in rabbit skeletal muscle: implications for regulation of actin-myosin interaction. Proceedings of the National Academy of Sciences of the United States of America 87:414–418.

56. Wolff, M. R., K. S. McDonald, and R. L. Moss. 1995. Rate of tension development in cardiac muscle varies with level of activator calcium. Circulation research 76:154–160.

57. Tesi, C., N. Piroddi, F. Colomo, and C. Poggesi. 2002. Relaxation kinetics following sudden Ca^2+^ reduction in single myofibrils from skeletal muscle. Biophys J 83:2142–2151.

58. Pate, E., K. Franks-Skiba, H. D. White, and R. Cooke. 1993. The use of differing nucleotides to investigate cross-bridge kinetics. J Biol Chem 268:10046–10053.

59. Potma, E. J., I. A. van Graas, and G. J. Stienen. 1995. Influence of inorganic phosphate and pH on ATP utilization in fast and slow skeletal muscle fibers. Biophys J 69:2580–2589.

60. Wang, L., A. Bahadir, and M. Kawai. 2015. High ionic strength depresses muscle contractility by decreasing both force per cross-bridge and the number of strongly attached cross-bridges. J Muscle Res Cell Motil 36:227–241.

61. Zhao, Y., P. M. Swamy, K. A. Humphries, and M. Kawai. 1996. The effect of partial extraction of troponin C on the elementary steps of the cross-bridge cycle in rabbit psoas muscle fibers. Biophys J 71:2759–2773.

62. Kentish, J. C. 1986. The effects of inorganic phosphate and creatine phosphate on force production in skinned muscles from rat ventricle. The Journal of physiology 370:585–604.

63. Cooke, R., K. Franks, G. B. Luciani, and E. Pate. 1988. The inhibition of rabbit skeletal muscle contraction by hydrogen ions and phosphate. J Physiol 395:77–97.

64. Cooke, R., and E. Pate. 1985. The effects of ADP and phosphate on the contraction of muscle fibers. Biophys. J. 48:789–798.

65. Reiser, P. J., and W. O. Kline. 1998. Electrophoretic separation and quantitation of cardiac myosin heavy chain isoforms in eight mammalian species. Am J Physiol 274:H1048–1053.

66. Pate, E., and R. Cooke. 1989. A model of crossbridge action: the effects of ATP, ADP and Pi. J Muscle Res Cell Motil 10:181–196.

67. Pate, E., K. Franks-Skiba, and R. Cooke. 1998. Depletion of phosphate in active muscle fibers probes actomyosin states within the powerstroke. Biophys J 74:369–380.

68. Cecchini, M., Y. Alexeev, and M. Karplus. 2010. Pi release from myosin: a simulation analysis of possible pathways. Structure (London, England : 1993) 18:458–470.

69. Yount, R. G., D. Lawson, and I. Rayment. 1995. Is myosin a “back door” enzyme? Biophys J 68:44S-47S; discussion 47S-49S.

70. Wang, L., and M. Kawai. 2013. A re-interpretation of the rate of tension redevelopment *k*TR in active muscle. J Muscle Res Cell Motil 34:407–415.

71. Brenner, B. 1991. Rapid dissociation and reassociation of actomyosin cross-bridges during force generation: a newly observed facet of cross-bridge action in muscle. Proc Natl Acad Sci USA 88:10490–10494.

72. Ebus, J. P., G. J. Stienen, and G. Elzinga. 1994. Influence of phosphate and pH on myofibrillar ATPase activity and force in skinned cardiac trabeculae from rat. The Journal of physiology 476:501–516.

73. Potma, E. J., and G. J. Stienen. 1996. Increase in ATP consumption during shortening in skinned fibres from rabbit psoas muscle: effects of inorganic phosphate. The Journal of physiology 496 (Pt 1):1–12.

74. Capitanio, M., M. Canepari, P. Cacciafesta, V. Lombardi, R. Cicchi, M. Maffei, … R. Bottinelli. 2006. Two independent mechanical events in the interaction cycle of skeletal muscle myosin with actin. Proc Natl Acad Sci USA 103:87–92.

75. Debold, E. P., S. Walcott, M. Woodward, and M. A. Turner. 2013. Direct observation of phosphate inhibiting the force-generating capacity of a miniensemble of Myosin molecules. Biophys J 105:2374–2384.

76. Linari, M., M. Caremani, and V. Lombardi. 2010. A kinetic model that explains the effect of inorganic phosphate on the mechanics and energetics of isometric contraction of fast skeletal muscle. Proc Biol Sci 277:19–27.

77. Mansson, A. 2019. The effects of inorganic phosphate on muscle force development and energetics: challenges in modelling related to experimental uncertainties. J Muscle Res Cell Motil. doi.org/10.1007/s10974-019-09558-2.

78. Amrute-Nayak, M., M. Antognozzi, T. Scholz, H. Kojima, and B. Brenner. 2008. Inorganic phosphate binds to the empty nucleotide binding pocket of conventional myosin II. J Biol Chem 283:3773–3781.

79. Poggesi, C., C. Tesi, and R. Stehle. 2005. Sarcomeric determinants of striated muscle relaxation kinetics. Pflugers Arch 449:505–517.

80. Herrmann, C., M. Houadjeto, F. Travers, and T. Barman. 1992. Early steps of the Mg^2+^-ATPase of relaxed myofibrils. A comparison with Ca^2+^-activated myofibrils and myosin subfragment 1. Biochemistry 31:8036–8042.

81. Stehle, R., C. Lionne, F. Travers, and T. Barman. 2000. Kinetics of the initial steps of rabbit psoas myofibrillar ATPases studied by tryptophan and pyrene fluorescence stopped-flow and rapid flow-quench. Evidence that cross-bridge detachment is slower than ATP binding. Biochemistry 39:7508–7520.

